# Mutation of the *Vinv* 5′ UTR regulatory region reduces acrylamide levels in processed potato to reach EU food-safety standards

**DOI:** 10.1101/2023.07.18.549223

**Authors:** Leonard Shumbe, Emanoella Soares, Yordan Muhovski, Inga Smit, Hervé Vanderschuren

## Abstract

Protein-coding regions of genes have so far been the preferred targets for trait improvement using CRISPR-Cas9 genome editing in plants. Alteration of the reading frame can result in termination of transcription or translation, hence, loss of function of the encoded protein. 5’ UTR sequences also represent a practical target region to alter gene expression and protein abundance. However, editing of 5’ UTRs has so far been scarcely used to engineer trait in crop plants. Here, we demonstrate that insertion of a single adenine nucleotide mediated by Cas9 in the 5’ UTR region of two alleles from the Vacuolar invertase gene (*VInv*) of the tetraploid potato variety Lady Rosetta (LaRo) is sufficient to substantially lower the content of reducing sugars in the potato under cold-storage conditions. Moreover, the acrylamide content generated during processing of the edited potato lines into crisps was more than three folds lower than the current EU-regulated maximum level of 750 μg/kg in crisps. This gene-editing approach represents a durable strategy to improve food safety of potatoes in varieties widely preferred by the consumers and the industry.

---

During several decades, the compound Chlorprofam (CIPC) was used as an efficient anti-sprouting agent for long term storage of potato. However, the recent prohibition of CIPC in the EU[1] is prompting the potato processing industry to search for alternative and safer anti-sprouting approaches[2]. In this context, storage at cold temperature (i.e. 4° C) has emerged as a valuable option for long term storage of potato without the use of CIPC. However, most commercial potato varieties processed by the industry accumulate high levels of reducing sugars during cold storage, a phenomenon called cold-induced sweetening (CIS). During high temperature processing of potatoes into products such as crisps and French fries, the reducing sugars react with free amino acids and peptides to produce the neurotoxin acrylamide, whose presence is evidenced by a brown-to-black coloration of the processed products[3]. Therefore it is key to prevent CIS in potato in order to unlock the potential of long-term storage at cold temperature in the processed potato value chain.

The potato-processing industry currently relies on only a few varieties that have been selected for their agronomical and organoleptic properties[4]. Because of the difficulty in breeding CIS-resistant potato varieties to replace the ones that are CIS-susceptible, New Genomic Techniques (NGTs) are emerging as useful approaches to rapidly introgress the CIS-resistant trait into commercial varieties used for processing into crisps and french fries.

Genome-editing techniques have revolutionised the field of plant breeding by enabling precise modification of crop genomes to rapidly add missing valuable traits in elite varieties. CRISPR/Cas9–based editing approaches have been instrumental to shorten the time for trait improvement in more complex tetraploid and hexaploid genomes. Despite the versatility of CRISPR-base approaches to target any selected sequence in a plant genome, the technology has so far been mainly used to target protein-coding sequences for trait improvement in crop plants[5–8].

The challenges faced by the EU potato processing industry are graphically depicted in Fig. 1. In the present work, we exploit editing of a non-coding 5’ UTR sequence as to engineer CIS resistance in an industry-preferred potato variety. Vacuolar invertase (VInv) has been identified as a key enzyme for conversion of sucrose into reducing sugars. Previous studies have demonstrated that silencing of the *VInv* gene is a suitable approach to lower the accumulation of reducing sugars upon cold storage of potato[3,8,9].

**Fig. 1.**
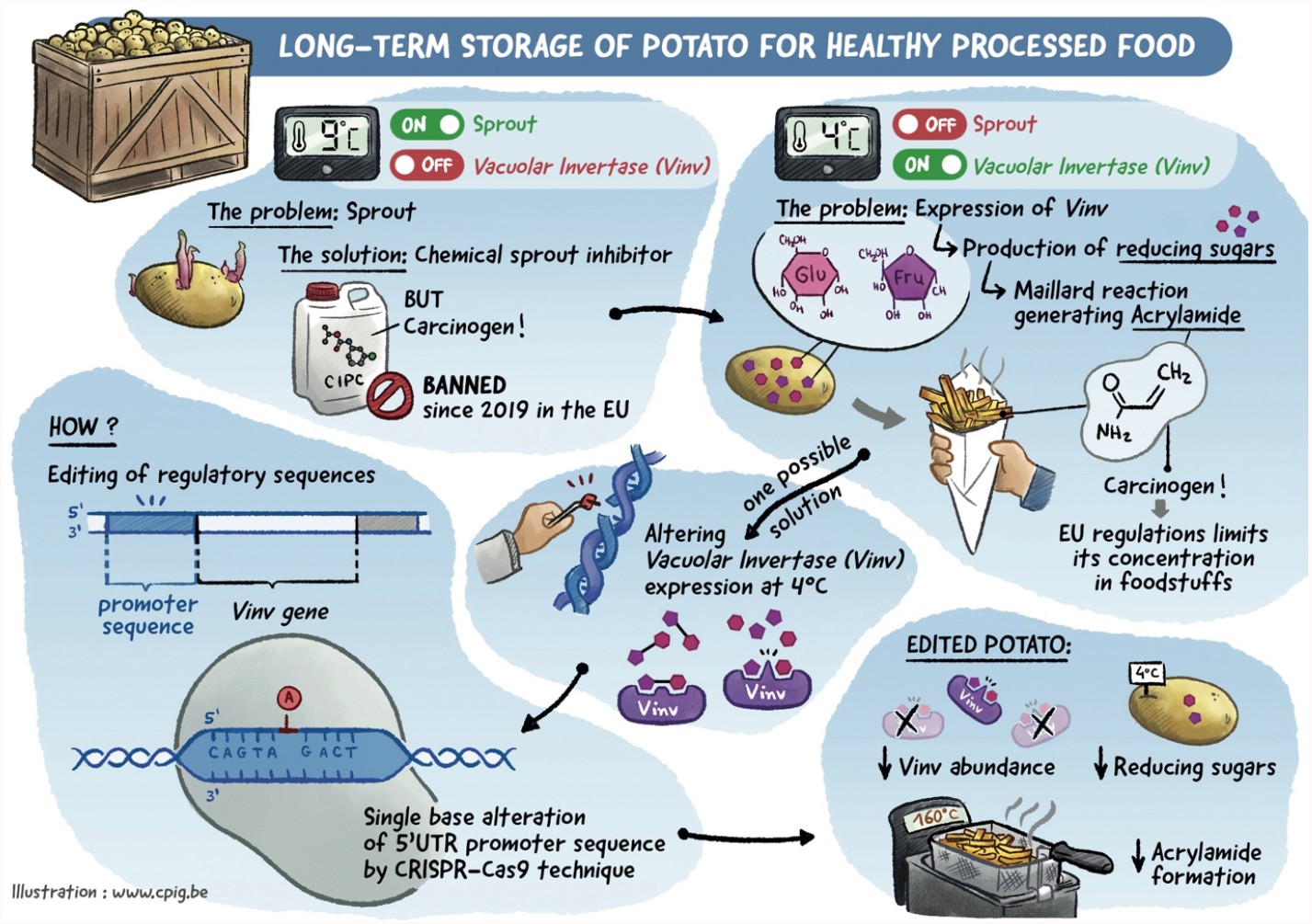
Graphical presentation of the challenges for long term storage of potato in the EU processing industry and the genome editing solution presented in the article.

We designed a short guide RNA (sgRNA) targeting the 5’ UTR region of the *VInv* gene using the CRISPR RGEN Cas-Designer Tool[10]. The sgRNA with the best score was selected to minimize off-target in the potato genome. Its activity was assessed in an *in vitro* cleavage assay using an amplicon from potato DNA as template (Fig.S1). We next transformed potato variety Lady Rosetta (LaRo) with the construct PC2300-sgRNA1–pcoCas9–eGFP, which contains a plant-codon-optimized *Cas9* sequence and the *Arabidopsis thaliana U6* promoter driving transcription of the sgRNA. LaRo is a variety used in the crisp industry and is CIS-susceptible[11]. Based on the sequence of the protospacer adjacent motif (PAM) recognized by the Cas9 used in the transformation (NGG), the selected sgRNA was expected to target two of the four alleles (50%) of LaRo *VInv* in their 5’ UTR sequences. Thirty-two independent potato lines were generated by *Agrobacterium*-mediated internode transformation, and were molecularly characterized by PCR–restriction enzyme (RE) assay on the genomic target sequence. Eleven of the thirty-two lines showed the desired genetic profile in the PCR–RE assay and their mini-tubers (T0 tubers) were vegetatively propagated to produce T1 tubers. T1 tubers of wild type (non-transformed) LaRo and Verdi varieties were also produced for use as controls. Verdi is a reference CIS-resistant potato variety^2^. Further analyses were performed on the T1 tubers.

We profiled the allelic variation in the 5’UTR target sequence from the T1 tubers by Illumina sequencing and identified four lines carrying one adenine insertion between positions -34 and -35 in the two editable alleles (P4, P6, P24 and P26), one transformed line without edits (P1), and six lines with different types and percentages of deletions in the two editable alleles (P5, P15, P21, P28, P30, P32) (Fig. 2A).

**Fig. 2.**
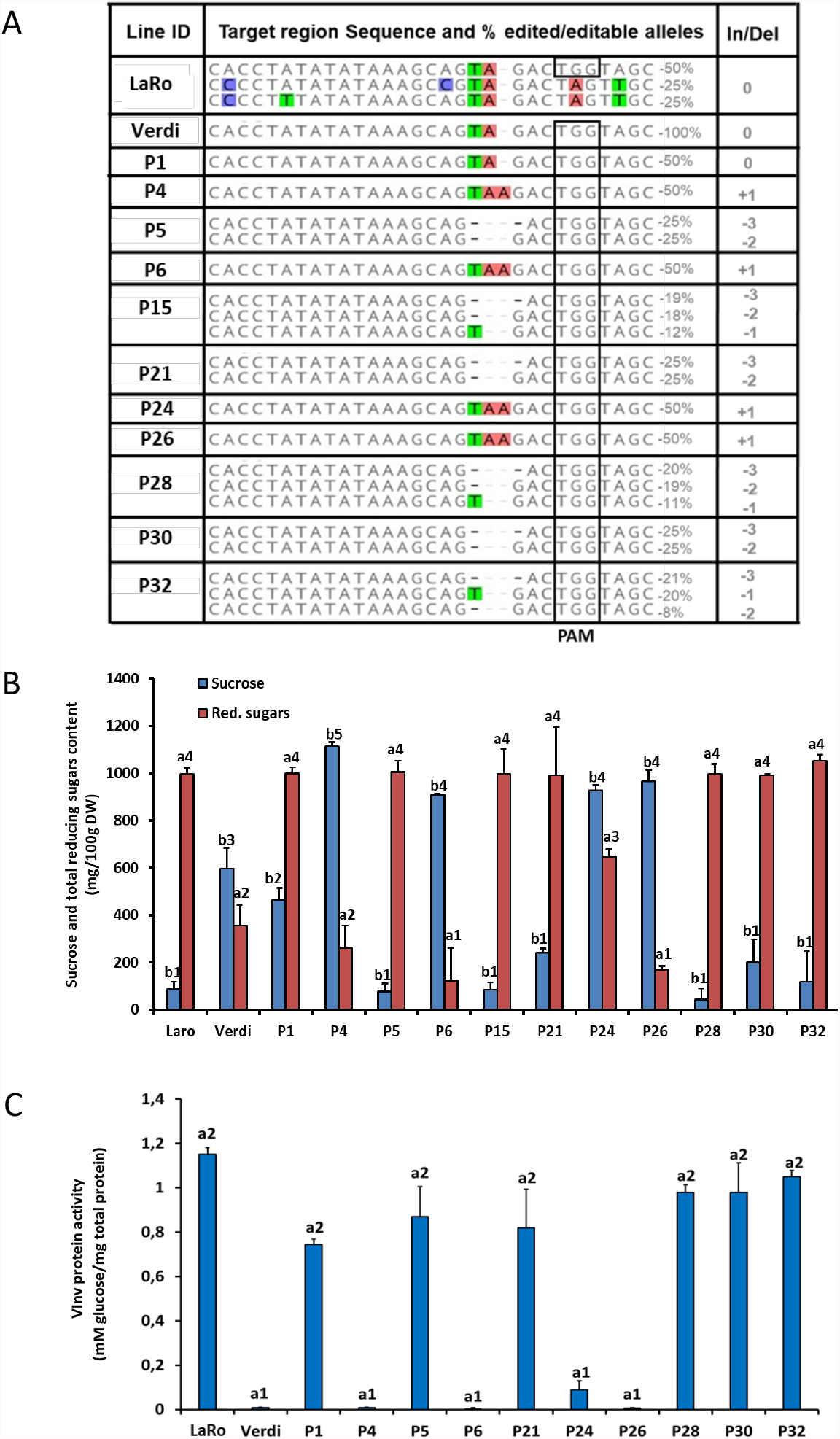
CIS-resistance engineered in Lady rosetta (LaRo) tubers. (**A**) target region from Illumina sequenced Lady Rosetta (LaRo) edited lines showing In-dels and the percentages of reads corresponding to the percentage of edited alleles. Wild-type Lady Rosetta and wild-type Verdi are non-transformed CIS-Susceptible and CIS-Resistant controls respectively. (**B**) Contents of sucrose and reducing sugars from tubers of non-transformed control, selected edited lines after 1 month storage at 4° C. (**C**) Activity of Vacuolar Invertase (VInv) quantified from potato tubers after 1 month of storage at 4°C. n=3 biological replicates for quantification of sugars and VInv protein activity. a1, a2, a3 and a4 indicate statistically significant differences in reducing sugars and VInv protein activity between the different lines and controls, and b1, b2, b3, b4, b5 represent statistically significant difference in sucrose content between the different lines and controls, at P<0.05 computed by Scott-Knott test.

The T1 tubers of the 11 lines and the control varieties were used to perform the first cold-storage assessment. Quantification of reducing sugars (glucose + fructose) in the selected 11 lines after storage at 4°C for 1 month revealed that the four lines with the single A-insertion (P4, P6, P24 and P26) contained significantly lower levels of reducing sugars compared with the non-edited control line (P1), and the other lines displaying various editing profiles of the 5’UTR (P5, P15, P21, P28, P30, P32) (Fig. 2B). Lines P6 and P26 had significantly lower levels of reducing sugars relative to the control CIS-resistant potato variety Verdi. There were no statistically significant differences in the levels of reducing sugars amongst the wild-type LaRo potatoes, the non-edited line (P1) and the edited lines P5, P15, P21, P28, P30, P32. In line with previous observations[3,12], sucrose contents in all samples inversely correlated with the contents of reducing sugars (Fig. 2B). These results indicate that the single A-insertion (P4, P6, P24 and P26) between positions -34 and -35 of the 5’ UTR of *VInv* gene was sufficient to render those lines CIS-resistant.

We assayed VInv enzymatic activity in selected lines, together with the wild-type LaRo and Verdi controls after storage at 4°C for 1 month. We observed significantly lower VInv activities in the lines with A-insertions and in the Verdi variety compared with the wild-type LaRo variety, the non-edited line and the lines with various editing profiles (Fig. 2C). The measured VInv activities were in line with the observed trends in reducing sugars (Fig. 2B), and the trends in transcript levels of *VInv* assayed in selected CIS-contracting T1 tubers after cold storage at 4°C for 1 month (Fig. S2A). These results indicate that a single A-insertion in 50% of the alleles at the 5’ UTR region cause a consistent and significant down-regulation of the *VInv* transcript and reduction in the levels of VInv activity upon cold storage.

We also assayed the levels of reducing sugars and sucrose in selected CIS-contrasting lines and varieties (P1, P4, P24, P26, P30, P32, LaRo and Verdi,) stored at room temperature for 1 month (Fig. S2B).

There was no statistically significant difference in the levels of reducing sugars between the CIS-susceptible lines (P1, P30, P32 and wild-type LaRo) and the CIS-resistant lines (P4, P24, P26 and Verdi). The sucrose levels in the CIS-resistant edited lines were similar to those detected in the CIS-resistant variety Verdi but were significantly lower than those detected in the CIS-susceptible lines and variety. This suggests that the single A-insertion in the 5’UTR also alters sucrose metabolism and accumulation in the tubers at room temperature — storage conditions widely used in households.

Furthermore, we assessed the effect of multi-generation vegetative propagation on the reducing sugars of selected CIS-contrasting lines and varieties (P1, P4, P24, P26, P30, P32, LaRo and Verdi). The levels of reducing sugars in the CIS-resistant lines were consistently significantly lower than the levels in the CIS-susceptible lines after two generations of propagation (T2 tubers) and after 1-month storage at 4°C (Fig.S2C). However, among the selected CIS-resistant lines, the levels of reducing sugars for line P26 were higher compared to the levels in the T1 tubers.

Finally, to assess the effect of the single A-insertion on the acrylamide contents of cold-stored potato upon high-temperature processing, T1 tubers of selected CIS-contrasting lines and their respective controls (Fig. 3A) were thin-sliced and fried in oil heated to 180°C for 3 minutes to produce crisps. The crisps were evaluated qualitatively by observing the differences in crisp color and quantitatively by measuring their acrylamide content with HPLC–MS. The fried crisps from CIS-susceptible lines P1, P21 and P28 and the non-transformed control LaRo variety displayed a brown-to-black color, while the ones from the CIS-resistant A-insertion lines (P4, P6 and P24) and the CIS-resistant variety Verdi consistently appeared pale yellow in color (Fig. 3B). The CIS-resistant edited lines showed acrylamide contents well below the benchmark value of 750 μg/kg set by the European Union for potato crisps[13]. Conversely the fried crisps from the CIS-susceptible lines and the wild-type variety LaRo had acrylamide contents above the EU benchmark value (Fig. 3C). These results also corroborate previous reports of a linear correlation between contents of reducing sugars, the color of crisps, and acrylamide content[3,11].

**Fig. 3.**
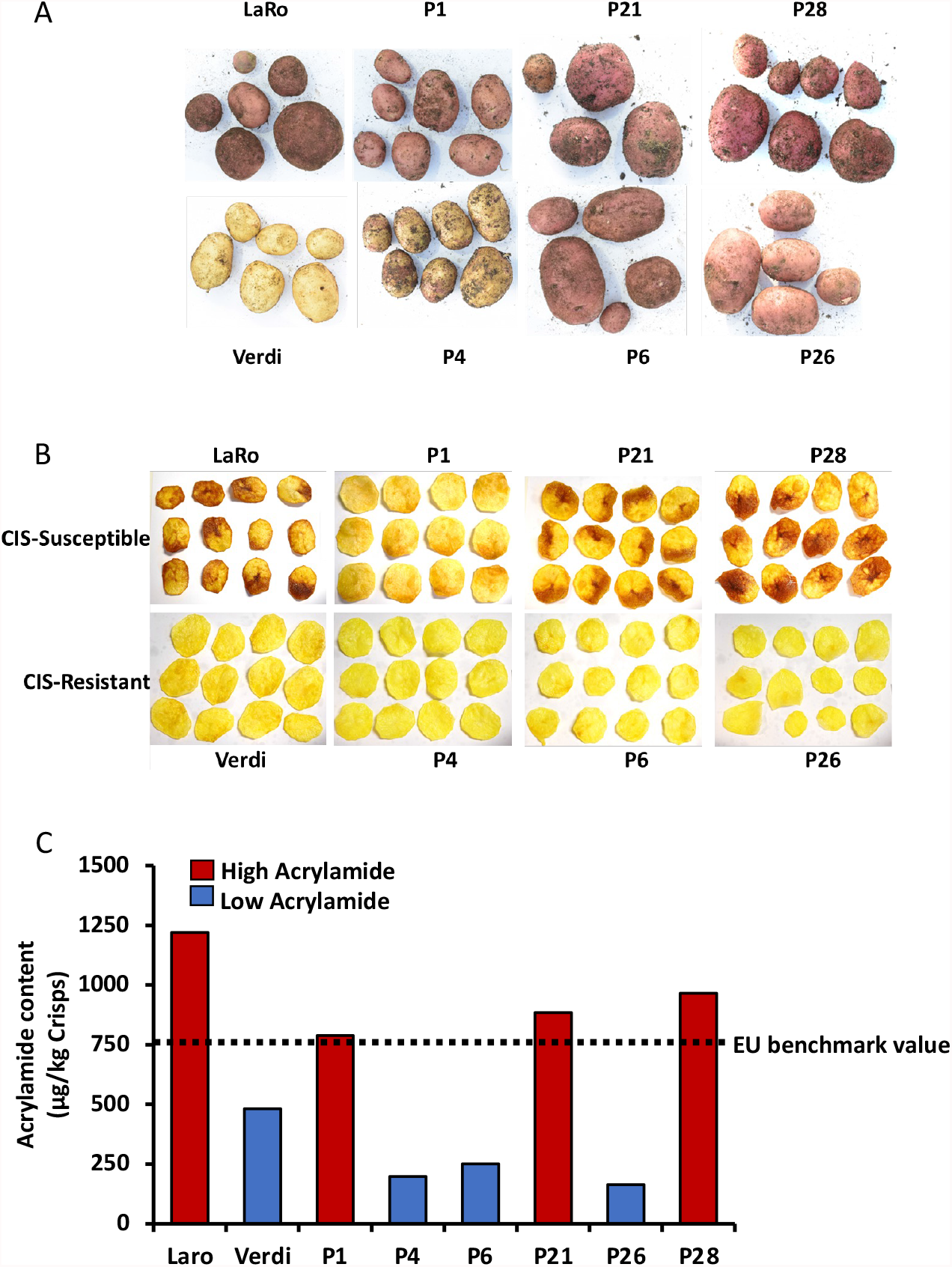
Tuber phenotype and Acrylamide content of CIS-contrasting lines and varieties. (**A**) Phenotype of tubers from edited lines shown to have contrasting levels of reducing sugars content after cold storage. (**B**) Crisps produced from CIS-contrasting tubers after cold-storage. (**C**) Acrylamide content of the crisps and indication of the EU benchmark level for acrylamide in crisps (n=2).

Taken together, our findings demonstrate that a single adenine insertion in the 5’UTR region of 50% of the LaRo *VInv* alleles is sufficient to alter the CIS phenotype of the LaRo variety, and thereby to reduce the acrylamide content in the processed products to an acceptable level within EU requirements. VInv is known to regulate several aspects of growth and development in plants[14,15], however, no abnormal phenotype was observed for the CIS-contrasting lines (Fig 3A), nor any significant differences in the mean tuber size (Fig. S2D).

Classical applications of gene-editing techniques for trait improvement in crop plants have preferentially targeted protein-coding sequences for double strand breaks and introduction of frame shift via the error prone non-homologous end joining (NHEJ) repair mechanism[16]. The 5’ UTR region has been shown to play a role in the regulation of mRNA stability[17] as it harbours the transcription initiation site in many eukaryotes. Therefore the 5’UTR region offers an alternative to the traditional exon targeting for genome editing[18-20]. Our data indicate that, by targeting 50% of the alleles in the 5’UTR of *VInv*, we quantitatively regulate gene expression and retain the desired phenotype of the tubers. While trait engineering usually relies on targeting all allelic coding sequences in potato[3,7,8], our study demonstrates that a stable CIS-resistance trait can be achieved by differential allele targeting of the *VInv* 5’ UTR. In some specific cases, differential allele targeting might also represent a more suitable approach to reduce the possible pleiotropic effects associated with silencing or knocking out of all alleles in polyploid crop species such as potato[9,21]. These results provide a timely solution to convert commercial CIS-susceptible potato varieties widely used by the European potato industry into CIS-resistant varieties for the subsequent production of safe and healthy crisps and French fries.

## Supplementary data

- Material and method
- Table S1, Figures S1and S2

## Acknowledgement

Authors acknowledge support from the EUREKA grant RESPRO (Région Wallonne – SPW6) and thank Jared Fudge for useful comments on the manuscript.

## Author Contribution

Conceptualization, LS and HV; methodology and experiments, LS, ES, YM and IS; formal analysis, LS, ES, YM and IS; resources, HV; writing—original draft preparation, LS; writing—review and editing, LS and HV; supervision, HV; project administration, HV; funding acquisition, HV.

