## Supplementary material for "Mutation of the *Vinv* 5′ UTR regulatory region reduces acrylamide levels in processed potato to reach EU food-safety standards": Material and Methods and Supplemental Figures

### Supplementary data

#### Material and methods

#### 1. sgRNA design and efficiency assay

The nucleotide sequence 5' -TATATATAAAGCAGTAGAC-3' located in the 5' UTR region of the potato *Vinv* gene was selected as the sgRNA target sequence, using the CRISPR RGEN Cas-Designer Tool[1]. The sequence was checked for off targets using the Cas-OFFinder tool[2] and by blasting against the potato genome on phytozome. The sgRNA cassette (T7promoter-target sequence-scaffold-terminator) was transcribed as described in the EnGen® sgRNA Synthesis Kit, *S. pyogenes* Protocol (NEB, USA).

The efficiency of the sgRNA was assessed in an *in vitro* cleavage assay, as described in the protocol for in vitro digestion of DNA with Cas9 Nuclease, S. pyogenes (NEB, USA). A 1.2 bk PCR amplicon spanning the sgRNA target sequence was used as the template in the cleavage reaction. The control reaction was performed following the same procedure without adding the *S. pyogenes* Cas9 nuclease.

#### 2. Editing and characterization of potato lines

##### 2.1 Generation of the transformation vector (PC2300-pcoCas9-AtU6-sgRNA)

The plant codon optimized Cas9 sequence (pcoCas9) was obtained from Addgene, in the PYPQ150 plasmid (Addgene Plasmid #69301). The CaMV-35S promoter was amplified from pCambia2300 plasmid using the primer pair ApaI-EcoRI-35s-F and NcoI-35s-R and cloned in the ApaI/NcoI sites of the pYPQ150 plasmid, to generate the pYPQ150_35S plasmid. The 35s-pcoCas9 fragment was then digested out from the pYPQ150_35s plasmid using EcoRI restriction enzyme and cloned into the EcoRI site of pCambia2300 to form pC2300-35S-pcoCas9 plasmid. The sgRNA cassette was assembled in a PCR reaction, using the primer pair BspEI-AtU6-gRNA-F and BspEI-Common-R and a synthesized sgRNA cassette as template. The purified sgRNA cassette was cloned into the PmeI site of the pC2300-35S-pcoCas9 plasmid, producing the pC2300-35S-pcoCas9-AtU6-sgRNA vector.

#### 2.2 Transformation of potato

Agrobacterium harboring the PC2300-pcoCas9-AtU6-sgRNA plasmid was grown in 20 mL LB liquid medium contain 20 mg/L of rifampicin, to OD600 of 0.9. The agro-suspension was centrifuged and resuspended in 10 mL of MS-liquid supplemented with acetosyringone to a final concentration of 100 µM. All steps in the transformation were performed under sterile conditions. Inter-nodal cuttings obtained from four weeks old in vitro LaRo potato plantlets were incubated with the agrobacterium harbouring the PC2300-pcoCas9-AtU6-sgRNA plasmid for 30 minutes at room temperature, after which the stem cuttings were blotted dry with sterile filter paper and placed on MS-solid medium (autoclaved 4.4 g/L MS with vitamin + 20 g/L sucrose + 3 g/L gelrite). After 2 days of co-cultivation at 28°C in the dark, excess agrobacterium on the stem cuttings were removed by washing twice with sterile distilled water and once with sterile distilled water supplemented with cefotaxime (300 mg/L). The cuttings were blotted to dry and place on Callus Induction Medium (CIM3: MS-solid medium supplemented with 3 mg/L BAP + 2.5 mg/L NAA + 300 mg/L cefotaxime +100 mg/L kanamycin). Once callus development was initiated (after about 10 days on CIM3), the cuttings were transferred to Shoot Induction Medium (SIM1: MS-solid medium supplemented with 2 mg/L BAP + 3 mg/L GA3 + 300 mg/L cefotaxime +100 mg/L kanamycin) for shoot production. Shoots of approximately 3 cm length were cut from distant points on a callus and from different calli and placed on MS-solid medium for root development. Once the roots had established, each plantlet was screened for the presence of the transgene, by PCR using the Phire Plant Direct PCR kit as described by the manufacturer (ThermoFisher Scientific, France) and primers targeting the kanamycin resistance gene (Kan500-F/R) and the Cas9 gene (Cas9-F/R). The plantlets positive for both kanamycin and Cas9 were grown and multiplied *in vitro* in MS-solid medium to generate at least four plantlets per line. Two plantlets per line were subsequently planted in soil to generate mini tubers (T0 tubers). The mini-tubers were stored at room temperature until dormancy was broken, then they were again planted in soil to produce tubers of caliber >30mm in diameter (T1 tubers). Tubers from selected lines were grown for a further generation to produce T2 tubers.

#### 2.3. Potato Phenotyping

The potato shoots were visually observed for any differences in growth between the transformed lines and the control varieties in all generations. The phenotype of the T1 generation tubers were assessed by visual examination for any abnormalities in the shapes of the tubers between the lines and control varieties, and by measuring the diameter of the four biggest tubers of each line and variety.

**2.4. Pre-screening for in-dels.**

DNA was extracted from the T0 tubers of all the lines generated, using the CTAB extraction method. The primer pair *vinv*1.2-F/R was used to PCR amplify a 1.2kb fragment spanning the target region of the gene. Restriction enzyme digestion with AccI was performed using 100 mg of each amplicon according to the manufacture's protocol (NEB, USA). Lines showing a range of digestion profiles (non-digested, partial digestion, full digestion) were selected for further analyses.

**2.5 Screening for in-dels**

DNA was extracted from a portion of the T1 tubers for each line using the CTAB extraction method. A 402 bp fragment was amplified from the DNA extracted from each line using the primer pair G1-adapt-F/R (containing the illumina adapter sequences at the 5'end of each primer) and Q5 polymerase (Bioke, Belgium). Each amplicon was sequenced using the 2*300 bp paired end MiSeq system (GIGA platform, University of Liège, Belgium), generating approximately 300,000 reads per sample. The sequencing data was analysed using the CRISPresso2 software[3], to identify in-dels in the different potato lines. The raw sequencing data is available at https://doi.org/10.5281/zenodo.8138764

**3. Sugar measurements**

D-Sucrose and total reducing sugars (D-Glucose and D-Fructose) were assayed from 200 mg of each freeze-dried, ground tuber, making use of the K-SUFRG kit (Megazyme, UK), as described in the assay procedure (Megazyme, K-SUFRG 04/17) with some modifications: 1 mL of distilled water was added to 200 mg of each freeze-dried powder sample in a 2 mL tube and vortexed until a homogenous suspension was obtained. The suspension was centrifuged at 15,000 rpm for 10 minutes at 4° C, and the supernatant was harvested for sugar quantification. D-sucrose, D-Glucose and D-Fructose content was quantified as described in the microplate procedure of the Megazyme K-SURFG assay procedure 04/17, using the Microplate 96 wells F-Bottom (Greiner bio-one, Germany). Each absorbance value was divided by the value 0.625 (a function of the diameter of the well and the total reaction volume), to adjust the path length of the microplate to 1 cm. Computation of content of the sugars in each sample was performed as described in the assay procedure.

**4. VInv activity assay**

VInv activity was assessed using the freeze-dried, ground, 4°C-stored potato samples of the T1 generation. Each sample was composed of 300 mg of pooled 5 biological replicates per line (60 mg per biological replicate). Four pooled replicates were per line were assessed. Total protein extraction, desaltingand enzyme assay was performed as previously described[4]. The glucose content of samples and controls was quantified using the K-SUFRG kit, as described in the assay procedure (Megazyme, K-SUFRG 04/17), and the amount of glucose produced was determined by calculating the difference in glucose content between samples and controls. Total protein content for each sample was quantified using the Bradford method[5] and vacuolar invertase activity was calculated as glucose concentration produced per hour per microgram of total protein.

**5. Acrylamide content**

A portion of the cold stored potato tubers from each line were processed into crisps by frying in oil at 180°C for 3 minutes. The crisps obtained from each line were later sealed in a plastic bag and manually ground into fine powder using a rolling pin. Acrylamide was extracted from the samples and quantified in chemical ionization mode by GC-MS on a Trace GC/Trace DSQ instrument (Thermo Scientific, Dreieich, Germany) as described previously[6]. Extraction and quantification were performed in duplicates.

**6. Gene expression**

*Vinv* gene expression was assayed from leaves and tubers by qRT-PCR. RNA extraction from tuber samples were performed as previously [7], with some modifications: Approximately 100 mg of lyophilized potato powder per sample was used and the volumes of the reagents were reduced five folds. RNA was extracted from fresh leaves using a protocol further modified from the protocol described for RNA extraction from tuber sample. Lysis was performed using RLT buffer (Qiagen, France). After centrifugation, equal volumes of chloroform:isoamyl alcohol (24:1) was added to the supernatant. No Tris-saturated phenol was used. The quality of the RNA from both tissues was assessed on a 1% agarose gel, after which DNAseI (Bioke, Netherlands) treatment was performed according to the manufacturer's instructions. cDNA was synthesized from 500 ng of each DNA-free RNA sample using GoTaq Reverse Transcription, Oligo dT kit (Promega, USA) according to the manufacturers' instructions. The cDNA was diluted 5 folds with distilled water and used in a reaction mix composed of 1x GoTaq qRT-PCR master mix (Promega, USA) and 10 nM primers for RT-qPCR. Each sample was analyzed in triplicates on a CFX96 Real-Time System (Bio-Rad, USA) using the following program: initial denaturation at 95° C for 3 mins, then 40 cycles of 95° C for 10 sec (denaturation), 60° C for 30 sec (Annealing) and 72° C for 30 sec (elongation), followed by a plate read. *18S rRNA was* used as the house keeping gene. The primer sequences for *Vinv* gene (*StVinv*-F/R*)* and the house keeping gene *(rRNA*-F/R*)* are listed in Table S1.

### 7. Statistical analyses

The students t-test and the Tukey test were used to compute the statistically significant difference between lines, varieties and between storage temperatures in this work.

**Table S1:** List of primers used in this work and their corresponding nucleotide sequences

| **Primer pair** | **Forward Primer sequence (5'-3')** | **Reverse Primer sequence (5'-3')** | **Size (bp)** |
|---|---|---|---|
| *Vinv* 1.2-F/R | AGCGAAGACAACAATGGTGA | CCGAGATGACAT-ACCCGAAT | 1200 |
| BspEI-AtU6-gRNA-F/ BspEI-Common-R | ACTTCCGGAGGAGTGATCAAAAGTCCCACATCGATCAGGTGATATATAGCAGCTTAGTTTATATAATGATAGAGTCGACATAGCGATTGTATATATAAAGCAGTAGACGTTTTAGAGCTAGAAATAGCAAGTT | ACTTCCGGAGCGTAATGCCAACTTTGTAC | 237 |
| ApaI-EcoRI-35s-F/ NcoI-35s-R | ACTGGGCCCGAATTCGTCAACATGGTGGAGCAC | ACTCCATGGGCGTGTCCTCTCCAAATG | 361 |
| Kan500-F/R | TGACTGGGCACAACAGACAA | CATGTGTCACGACGA-GATCCT | 500 |
| Cas9-F/R | ACACACTTATAAACTACAGAAAAGCA | GGCTCTTGTTAGACAG-CAGC | |
| G1-adapt-F/R | TCGTCGGCAGCGTCAGATGTGTATAAGAGACAGAC-CAAACTAGCCTAAGGACCA | GTCTCGTGGGCTCGGA-GATGTGTATAAGAGACAG-CATGCAG-CAATGCTCATGGG | 402 |
| St*Vinv*-F/R | GGGTATGTGGGAGTGTGTGG | ATTCCACAATCCAATTCCGGGT | 201 |
| 18S *rRNA*-F/R | GGGCATTCGTATTTCATAGTCAGAG | CGGTTCTTGAT-TAATGAAAACATCCT | 101 |

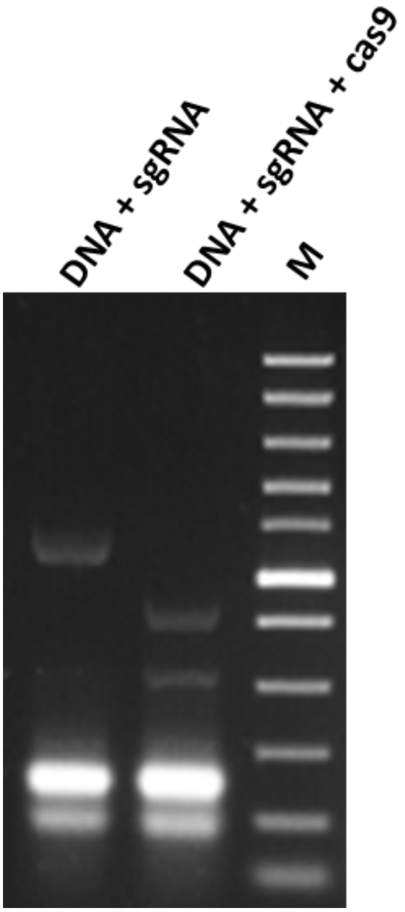


**Fig. S1**: *In vitro cleavage activity of target sgRNA on PCR-amplified DNA from potato. The lowest two bands in the first and second lanes represent monomers and dimers of the sgRNA respectively.*

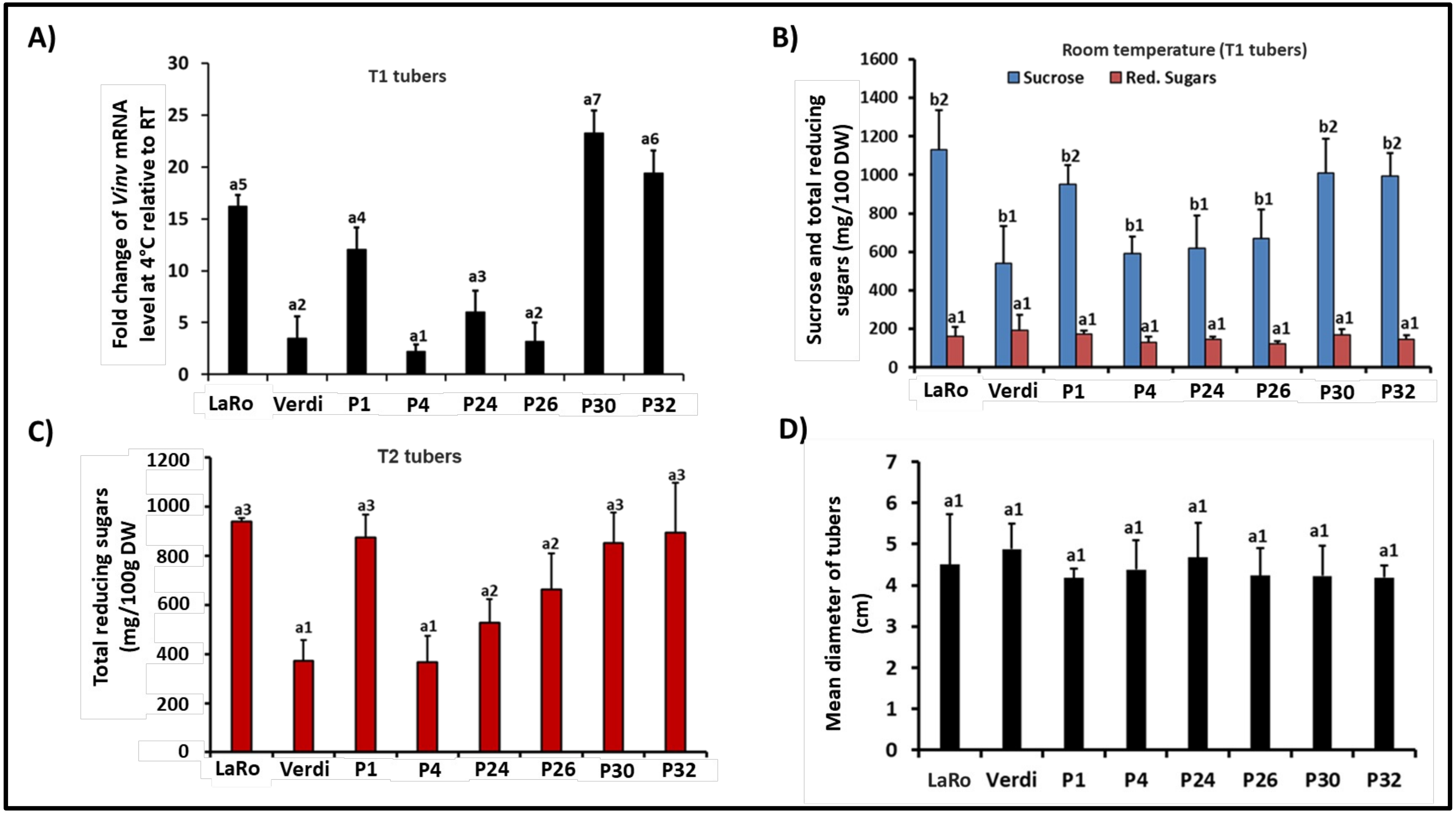


**Fig. S2:** *Characterization of selected CIS-contrasting lines and varieties. (**A**) Fold change of VInv gene expression in tubers stored for 1 month at 4° C relative to tubers stored at room temperature for the same duration. (**B**) Sucrose and reducing sugars content quantified from CIS-contrasting potato lines and controls after storage for 1 month at room temperature. (**C**) Reducing sugars quantified from CIS-contrasting lines multiplied after 3 generations (T2) and stored at 4° C for 1 month (**D**) mean diameter of four largest tubers from selected T1 generation lines and varieties. a1, a2, a3… a7 indicate statistically significant differences in reducing sugars and VInv gene expression between the different lines and controls, and b1, b2 represent statistically significant difference in sucrose content between the different lines and controls, at P<0.05 computed by Scott-Knott test.*
